# A translational rodent model of individual differences in sensitivity to the aversive properties of ethanol

**DOI:** 10.1101/2023.06.08.544209

**Authors:** Kathryn R Przybysz, Lindsey A Ramirez, Joseph R Pitock, E Margaret Starr, Hyerim Yang, Elizabeth J Glover

## Abstract

**Background:** A strong relationship exists between individual sensitivity to the aversive properties of ethanol and risk for alcohol use disorder (AUD). Despite this, our understanding of the neurobiological mechanisms underlying subjective response to ethanol is relatively poor. A major contributor to this is the absence of preclinical models that enable exploration of this individual variability similar to studies performed in humans.

**Methods:** Adult male and female Long-Evans rats were trained to associate a novel tastant (saccharin) with acute exposure to either saline or ethanol (1.5 g/kg or 2.0 g/kg i.p.) over three conditioning days using a standard conditioned taste aversion (CTA) procedure. Variability in sensitivity to ethanol-induced CTA was phenotypically characterized using a median split across the populations studied.

**Results:** When examining group averages, both male and female rats that had saccharin paired with either dose of ethanol exhibited reduced saccharin intake relative to saline controls of ethanol-induced CTA. Examination of individual data revealed a bimodal distribution of responses uncovering two distinct phenotypes present in both sexes. CTA-sensitive rats exhibited a rapid and progressive reduction in saccharin intake with each successive ethanol pairing. In contrast, saccharin intake was unchanged or maintained after an initial decrease from baseline levels in CTA-resistant rats. While CTA magnitude was similar between male and female CTA-sensitive rats, CTA-resistant females were more resistant to the development of ethanol-induced CTA than their male counterparts. Phenotypic differences were not driven by differences in baseline saccharin intake. CTA sensitivity correlated with behavioral signs of intoxication in only a subset of rats.

**Conclusions:** These data parallel work in humans by revealing individual differences in sensitivity to the aversive properties of ethanol that emerge immediately after initial exposure to ethanol in both sexes. This model can be leveraged in future studies to investigate the neurobiological mechanisms that confer risk for AUD.

## Introduction

Heavy alcohol use can lead to a diagnosis of alcohol use disorder (AUD), which poses a significant health risk to affected individuals (Griswold et al., 2018; *Key Substance Use and Mental Health Indicators in the United States: Results from the 2020 National Survey on Drug Use and Health*, 2020). While much of the research aimed at understanding the neurobiological mechanisms underlying vulnerability for AUD has focused on the rewarding properties of ethanol, its aversive properties also affect the decision to drink (Riley, 2011; Verendeev & Riley, 2013). Indeed, while rewarding properties like euphoria, anxiolysis, and social facilitation, can promote continued drinking, ethanol’s aversive properties, which include sedation, motor impairment, and dysphoria, can limit drinking (A. C. King et al., 2019a; Riley, 2011; Verendeev & Riley, 2013). Importantly, individuals differ in their sensitivity to these properties, leading to individual variability in the subjective response to ethanol (Holdstock et al., 2000; A. C. King et al., 2002; Morean & Corbin, 2010). This subjective response is a significant risk factor for the development of AUD (Hines et al., 2005; Schuckit, 1999), such that individuals who are more sensitive to the rewarding properties of ethanol and less sensitive to its aversive properties are more likely to drink heavily (Holdstock et al., 2000; A. C. King et al., 2002, 2011; Verendeev & Riley, 2013). In fact, lower sensitivity to the aversive properties of ethanol is predictive of future binge drinking and AUD diagnosis (A. C. King et al., 2011, 2014).

Preclinical models using either ethanol-induced conditioned taste aversion (CTA) or conditioned place aversion (CPA) to assess response to the aversive properties of ethanol have successfully recapitulated the aforementioned clinical findings. For example, rat strains selectively bred for high ethanol drinking or preference exhibit less ethanol-induced CTA compared to their low drinking/preferring counterparts or their founder strains (Barkley-Levenson et al., 2015; Brunetti et al., 2002; Crabbe et al., 2019; Dyr et al., 2016; Wyszogrodzka et al., 2021). Similarly, DBA mice are more sensitive to ethanol-induced CTA than C57 mice, which typically drink higher quantities of ethanol than the DBA strain (Risinger & Cunningham, 1992). Studies assessing aversion to ethanol across multiple inbred mouse strains have also shown that stronger ethanol-induced CTA (Broadbent et al., 2002; Phillips et al., 2005) or CPA (Cunningham, 2019) was significantly correlated with lower ethanol intake or preference. Furthermore, a meta-analysis summarizing findings across 182 studies found that mouse strains with greater home-cage, voluntary ethanol intake exhibited lower ethanol-induced CTA or CPA (Green & Grahame, 2008). A more recent review drew similar conclusions, lending further support to the negative relationship between ethanol-induced CTA/CPA and drinking (Seemiller et al., 2022).

In addition to findings from inbred rodent strains, a number of studies have also shown differences in sensitivity to the aversive properties of ethanol across sexes and throughout development. For example, adolescent rodents are less sensitive to ethanol-induced CTA than adults, which likely contributes to greater levels of binge drinking in this population (Saalfield & Spear, 2016; Schramm-Sapyta et al., 2010, 2014; Vetter-O’Hagen et al., 2009a). Similarly, female rats, which typically consume greater quantities of ethanol than males, are less sensitive to ethanol’s aversive properties than males (Cailhol & Mormède, 2002; Roma et al., 2006, 2007; Schramm-Sapyta et al., 2014; Vetter-O’Hagen et al., 2009a) although this appears to vary by age (Schramm-Sapyta et al., 2014; Vetter-O’Hagen et al., 2009a) and strain (Cailhol & Mormède, 2002).

While these studies provide insight into the potential genetic and demographic factors contributing to risk for AUD, to our knowledge, no studies have examined individual differences in sensitivity to the aversive properties of ethanol in outbred animals of the same age or sex. Such comparisons are likely to have greater translational relevance to assessments done in the clinical population. To address this gap, the present study examined individual differences in ethanol-induced CTA in adult male and female outbred Long-Evans rats. Our results reveal that sensitivity to ethanol-induced CTA is bimodally distributed in both males and females, with some individuals exhibiting relatively high sensitivity and others relative resistance to CTA. The high degree of variability in sensitivity to the aversive properties of ethanol observed in this population parallels the individual differences in subjective response to ethanol observed in humans.

## Materials & Methods

### Animals

Adult male and female Long-Evans rats (Envigo, Indianapolis, IN) were P60 on arrival and allowed to acclimate to the vivarium for at least one week prior to experimental procedures. All rats were singly housed in standard cages in a temperature-controlled room on a reverse 12:12h light/dark cycle (lights on at 22:00). Food (Teklad 7912, Envigo) was provided *ad libitum* throughout the duration of the experiment. Water was available *ad libitum* prior to the onset of CTA procedures (see below). All procedures were approved by the University of Illinois Chicago Institutional Animal Care and Use Committee and adhered to the NIH Guidelines for the Care and Use of Laboratory Animals.

### Conditioned taste aversion

Conditioned taste aversion was used to assess sensitivity to the aversive properties of ethanol using procedures described previously by our laboratory (Glover et al., 2016) (**Figure 1A**). In brief, rats were habituated to scheduled access to water for 30 min/day for seven days after which they were conditioned to associate a novel 0.1% saccharin solution with an intraperitoneal (i.p.) injection of either ethanol (EtOH, 1.5 g/kg or 2.0 g/kg) or saline (EtOH equivalent volume, n=24-26/sex/drug group). Conditioning days were separated by three water recovery days. Bottle weights were taken before and after each session to measure fluid consumption during the 30-min access period. Rats receiving ethanol were assessed for behavioral signs of intoxication using a previously published subjective rating scale with scores ranging from 1 (no signs of intoxication) to 5 (loss of consciousness) (Glover et al., 2019; Trantham-Davidson et al., 2014).

**Figure 1.**
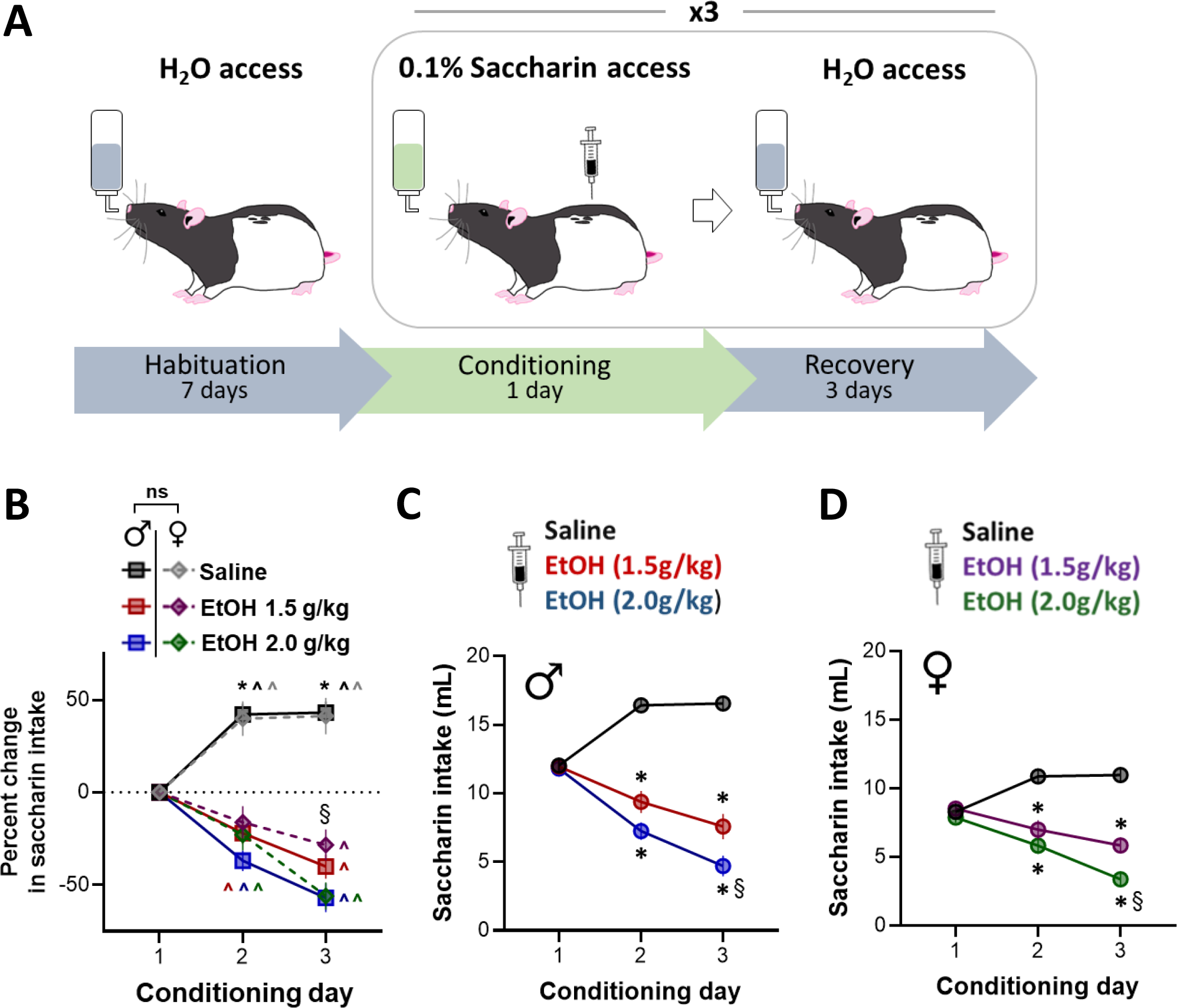
Dose-dependent increase in ethanol-induced CTA in adult male and female Long-Evans rats. **(A)** Rats underwent three conditioning sessions during which 0.1% saccharin was followed by injection of either saline, 1.5 g/kg EtOH, or 2.0 g/kg EtOH. Each conditioning day was separated by three water recovery days. **(B)** Change in saccharin intake in males and females normalized to baseline as percent change revealed a significant difference in intake between saline and 1.5 g/kg and 2.0 g/kg EtOH groups on conditioning days 2 and 3. Intake was also significantly different between the 1.5 g/kg and 2.0 g/kg EtOH groups on conditioning day 3. Significant changes in intake from baseline were also observed on conditioning days 2 and 3 for rats within each sex/drug group. Both male **(C)** and female **(D)** rats injected with 1.5 g/kg or 2.0 g/kg EtOH drank significantly less saccharin than saline-injected rats on conditioning days 2 and 3. Rats injected with 2.0 g/kg EtOH also drank significantly less saccharin than rats injected with 1.5 g/kg EtOH on conditioning day 3. *p<0.0001 saline compared to both EtOH doses; ^§^p≤0.05 compared to 1.5 g/kg EtOH; ^p<0.05 compared to conditioning day 1.

### Phenotypic classification

Rats were categorized into CTA-sensitive and -resistant phenotypes upon completion of the study using a median split of the percent change in saccharin intake from baseline (conditioning day 1, prior to drug injection) to conditioning day 3. Rats with a percent change in intake that was numerically below the median, indicating greater magnitude CTA, were classified as CTA sensitive. Conversely, rats with a percent change in intake that was numerically above the median, indicating lower magnitude CTA, were classified as CTA resistant.

### Statistical analysis

Saccharin intake and comparisons between phenotypes were analyzed using multifactorial analysis of variance (ANOVA), unpaired t-tests, and one-sample t-tests as described below. Simple linear regressions were used to assess the relationship between intoxication score and percent change from baseline saccharin intake. Where appropriate, *post-hoc* analyses were performed using Tukey’s HSD test. ROUT outlier tests were run on all parameters. One female from the 1.5 g/kg EtOH group and one female from the 2.0 g/kg EtOH group were identified as statistical outliers and removed from all subsequent analyses. Data from one additional female rat in the saline group was removed due to a technical error resulting in inaccurate measurement of fluid consumption. All statistical tests and graphs were generated using Prism 9 (Graphpad Software, San Diego, CA) or SPSS (IBM, New York). Data are reported as mean ± SEM.

## Results

### Ethanol induces significant conditioned taste aversion in males and females

To determine the extent of ethanol-induced CTA at each of the two doses administered, we conducted an omnibus ANOVA on saccharin intake with conditioning day, drug group, and sex as factors. This analysis uncovered a significant main effect of sex [*F*(1, 146)=70.650, *p*<0.0001], which showed that, overall, males consumed significantly more saccharin than females independent of drug group. Therefore, to better determine whether sex differences in CTA magnitude were present, raw intake data were normalized to percent change in intake from baseline (**Figure 1B**). Using this approach, there were significant main effects of conditioning day [*F*(1, 146)=26.31, *p*<0.001] and drug [*F*(2, 146)=85.61, *p*<0.001], but not sex [*p*>0.05]. There was also a significant conditioning day x drug interaction [*F*(4, 292)=56.91, *p*<0.001]. Post-hoc analyses revealed a significant difference between saline and 1.5 g/kg EtOH on conditioning days 2 [*p*<0.001] and 3 [*p*<0.001], as well as a significant difference between saline and 2.0 g/kg EtOH on conditioning days 2 [*p*<0.001] and 3 [*p*<0.001]. Additionally, a significant difference between 1.5 g/kg EtOH and 2.0 g/kg EtOH was present on conditioning day 3 [*p*=0.014]. To more directly assess changes in saccharin intake from baseline across conditioning days within each drug group and sex, we also performed one-sample t-tests on conditioning days 2 and 3 in these groups. In males on conditioning day 2, there was a significant increase from baseline in the saline group [*t*(25)=7.11, *p*<0.001] and a significant decrease from baseline in the 1.5 g/kg EtOH group [*t*(25)=3.83, *p*=0.0008] and the 2.0 g/kg EtOH group [*t*(25)=6.59, *p*<0.0001]. Similarly, on conditioning day 3, there was also a significant increase from baseline in the saline group [*t*(25)=7.34, *p*<0.0001] and a significant decrease from baseline in the 1.5 g/kg EtOH group [*t*[25)=6.07, *p*<0.0001] and the 2.0 g/kg EtOH group [*t*(25)=7.33, *p*<0.0001]. In females on conditioning day 2, we found a significant increase from baseline in the saline group [*t*(24)=4.26, *p*=0.0003], a trend toward a decrease from baseline in the 1.5 g/kg EtOH group [*t*(23)=1.86, *p*=0.07], and a significant decrease from baseline in the 2.0 g/kg EtOH group [*t*(24)=2.69, *p*=0.01]. On conditioning day 3 in females, we again found a significant increase from baseline in the saline group [*t*(24)=4.18, *p*=0.0003], as well as a significant decrease from baseline in both the 1.5 g/kg EtOH group [*t*(23)=3.39, *p*=0.0025] and the 2.0 g/kg EtOH group [*t*(24)=7.59, *p*<0.0001].

Similar results were obtained when using a two-way repeated measures mixed ANOVA with conditioning day and drug as factors to analyze changes in raw intake separately for males and females. Thus, in males, this analysis revealed a significant main effect of conditioning day [*F*(1.86, 139.5)=17.86, *p*<0.0001], a significant main effect of drug [*F*(2, 75)=47.29, *p*<0.0001], and a significant conditioning day x drug interaction [*F*(4, 150)=44.55, *p*<0.0001]. Post-hoc analyses showed significantly lower saccharin intake in the 1.5 g/kg EtOH group compared to saline on conditioning days 2 [*q*(41.42)=10.53, *p*<0.0001] and 3 [*q*(36.55)=12.26, p<0.0001]. This was also true for the 2.0 g/kg EtOH group relative to saline controls at the same time points [day 2: *q*(46.80)=15.97, *p*<0.0001; day 3: *q*(41.36]=18.94, *p*<0.0001]. Saccharin intake was also significantly lower on conditioning day 3 in the 2.0 g/kg EtOH group compared to intake in the 1.5 g/kg EtOH group on the same day [*q*(48.04)=3.397, *p*≤0.05] (***Figure 1C***).

In females, analysis of saccharin intake across days revealed significant main effects of conditioning day [*F*(1.876, 133.2)=10.51, p<0.0001], and drug [*F*(2, 71)=34.18, *p*<0.0001], as well as a significant conditioning day x drug interaction [*F*(4, 142)=20.78, *p*<0.0001]. Post-hoc analyses revealed findings similar to those observed in the males. Specifically, rats that had 1.5 g/kg EtOH paired with saccharin had significantly lower saccharin intake compared to saline controls on conditioning days 2 [*q*(38.48)=6.761, *p*<0.0001] and 3 [*q*(45, 49)=9.462, *p*<0.0001]. Similarly, the 2.0 g/kg EtOH group exhibited significantly lower saccharin intake compared to saline on both conditioning days [day 2: *q*(43.21)=9.664, *p*<0.0001; day 3: *q*(47.46)=14.46, *p*<0.0001]. The 2.0 g/kg EtOH group also had significantly lower saccharin intake compared to the 1.5 g/kg EtOH group on conditional day 3 [*q*(46.73)=4.339, *p*=0.001] (***Figure 1D***). Taken together, these data demonstrate the development of dose-dependent ethanol-induced CTA of similar magnitude between male and female rats.

### Individual differences in sensitivity to ethanol-induced CTA

To determine whether individual differences in sensitivity to the aversive properties of ethanol could be categorized into distinct phenotypes, we calculated the percent change in saccharin intake from baseline, prior to any ethanol exposure, to intake on conditioning day 3, after two saccharin-ethanol pairings. This analysis revealed a bimodal distribution of responses present in both males (**Figure 2A,2D**) and females (**Figure 3A,3D**) at both doses tested. For one cluster of rats, the percent change in saccharin intake was relatively close to zero, indicating minimal reduction in consumption across conditioning days. In the other cluster of rats, the percent change in saccharin intake was substantially below zero, indicating a large reduction in consumption across conditioning days. We refer to these clusters as CTA-resistant and -sensitive, respectively. Importantly, saccharin intake did not differ significantly at baseline between phenotypes. This was evident in male rats injected with 1.5 g/kg EtOH using an unpaired t-test comparing saccharin intake on conditioning day 1 between rats classified as CTA resistant and CTA sensitive [*t*(16.46)=1.95, *p*>0.05] (***Figure 2B***). Likewise, no difference in baseline saccharin intake was observed between phenotypes in males injected with 2.0 g/kg EtOH [*t*(24)=0.467, *p*>0.05] (***Figure 2E***). Similar to males, we observed no significant difference in baseline saccharin intake between phenotypes in females at either dose [1.5 g/kg EtOH: *t*(22)=1.282, *p*>0.05; 2.0 g/kg EtOH: *t*(23)=0.205, *p*>0.05] (**Figures 3B, *3E***). Together, these data suggest that sensitivity to ethanol-induced CTA is not driven by innate differences in hedonic drive for saccharin in either males or females.

**Figure 2.**
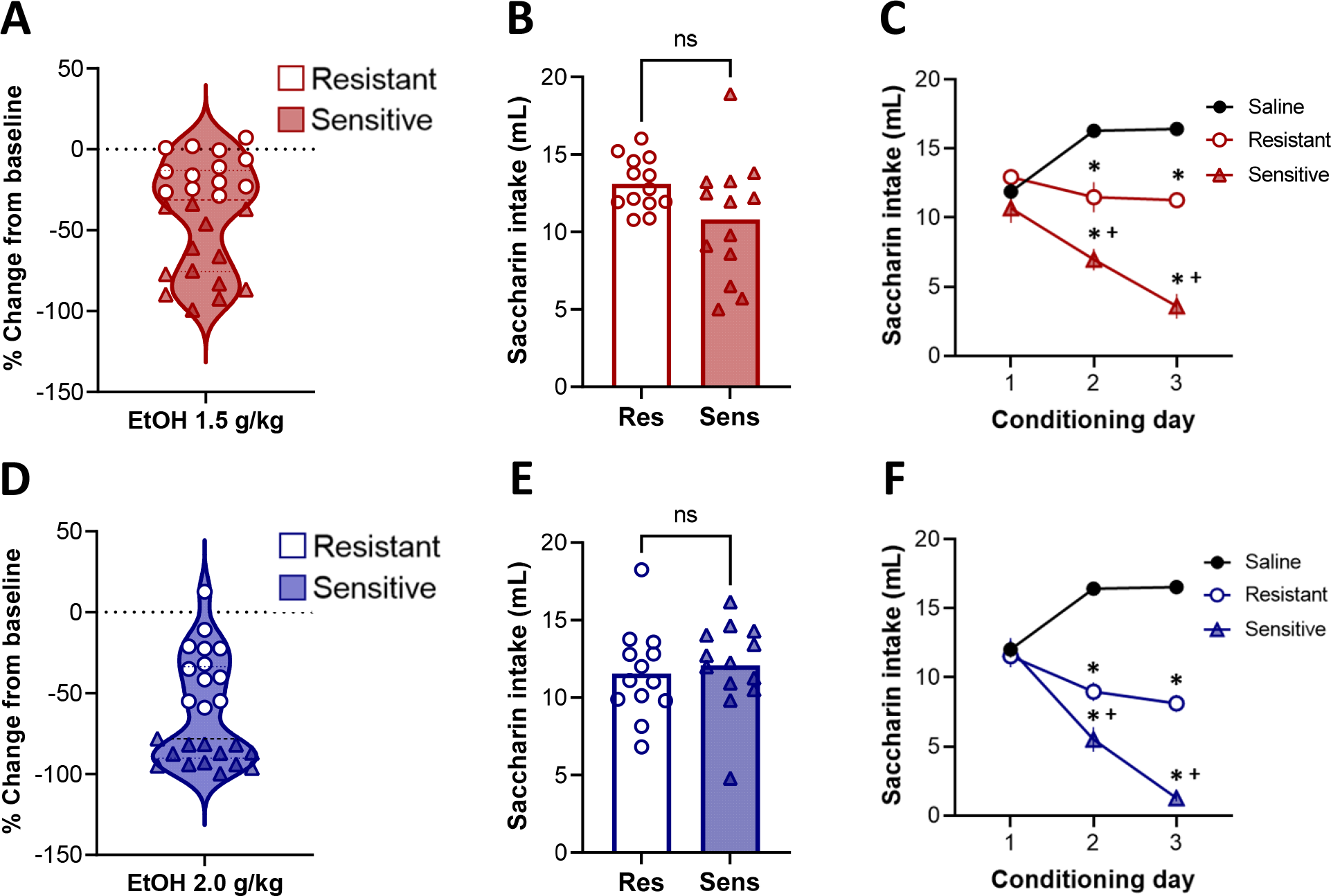
Individual differences in sensitivity to ethanol-induced CTA in male Long-Evans rats. A bimodal distribution in sensitivity to ethanol-induced CTA is apparent in rats injected with either 1.5 g/kg **(A)** or 2.0 g/kg **(D)** EtOH when individual data is plotted as percent change from baseline in saccharin intake on conditioning day 3. Rats were classified as CTA sensitive or CTA resistant using a median split. Baseline saccharin intake (prior to pairing with EtOH) was not significantly different between CTA-resistant and -sensitive rats in either the 1.5 g/kg EtOH group **(B)** or the 2.0 g/kg EtOH group **(E)**. Average saccharin intake differed between phenotypes across conditioning days. In both the 1.5 g/kg EtOH group **(C)** and the 2.0 g/kg EtOH group **(F)**, both CTA-resistant and -sensitive rats drank significantly less saccharin than saline controls on conditioning days 2 and 3. CTA-sensitive rats in both drug groups also drank significantly less saccharin than CTA-resistant rats on conditioning days 2 and 3. **\***p<0.002 compared to saline; ^+^p<0.01 compared to CTA-resistant. In A and D, dashed lines indicate the median and dotted lines indicate the upper and lower quartiles.

**Figure 3.**
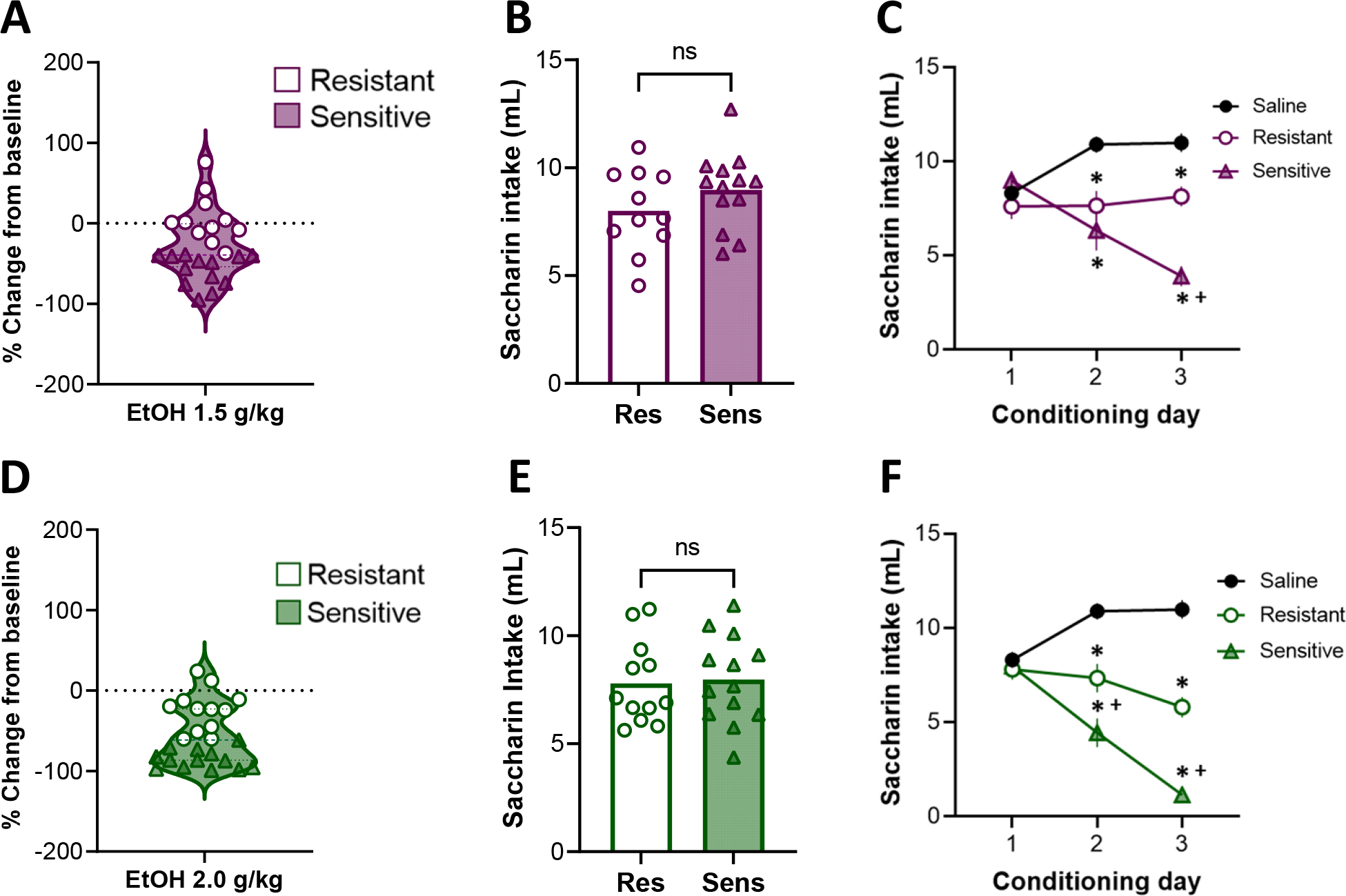
Individual differences in sensitivity to ethanol-induced CTA in female Long-Evans rats. Similar to males, a bimodal distribution in sensitivity to ethanol-induced CTA is apparent in rats injected with either 1.5 g/kg **(A)** or 2.0 g/kg **(D)** EtOH when individual data is plotted as percent change from baseline in saccharin intake on conditioning day 3. Rats were classified as CTA-sensitive or CTA-resistant using a median split. Baseline saccharin intake (prior to pairing with EtOH) was not significantly different between CTA-resistant and -sensitive rats in either the 1.5 g/kg EtOH group **(B)** or the 2.0 g/kg EtOH group **(E)**. Average saccharin intake differed between phenotypes across conditioning days. In both the 1.5 g/kg EtOH group **(C)** and the 2.0 g/kg EtOH group **(F)**, both CTA-resistant and -sensitive rats drank significantly less saccharin than saline controls on conditioning days 2 and 3. CTA-sensitive rats injected with 1.5 g/kg EtOH (C) also drank significantly less saccharin than CTA-resistant rats on conditioning day 3. In contrast, the difference in intake between CTA-sensitive and -resistant phenotypes injected with 2.0 g/kg EtOH (F) was significant on conditioning days 2 and 3. **\***p<0.002 compared to saline; ^+^p<0.01 compared to CTA-resistant. In A and D, dashed lines indicate the median and dotted lines indicate the upper and lower quartiles.

To determine whether the emergence of ethanol-induced CTA was significantly different between phenotypes, we next analyzed saccharin intake across conditioning days by comparing rats that received EtOH to saline controls. In males that received saccharin paired with 1.5 g/kg EtOH, a two-way mixed ANOVA revealed significant main effects of conditioning day [*F*(1.90, 93.19=6.933, *p*=0.002], and of phenotype [*F*(2, 49)=49.47, *p*<0.0001] and a conditioning day x phenotype interaction [*F*(4, 98= 43.9, *p*<0.0001] (**Figure 2C**). Post-hoc analyses showed that, while there were no baseline differences in saccharin intake between CTA-resistant or -sensitive rats compared to saline-injected rats, intake diverged significantly between groups on later conditioning days. Both CTA-sensitive and -resistant rats consumed significantly less saccharin compared to the saline group on conditioning days 2 [sensitive: *q*(21.6)=14.28, *p*<0.0001; resistant: *q*(36.49)=2.062, *p*=0.002] and 3 [sensitive: *q*(18.62)=17.97, *p*<0.0001; resistant: *q*(30.29)=10.61, *p*<0.0001]. CTA-sensitive rats also had significantly lower saccharin intake compared to CTA-resistant rats on conditioning days 2 [q(21.74)=4.74, p=0.008] and 3 [*q*(18.95)=10.54, *p*<0.0001].

Similar results were uncovered when comparing rats injected with 2.0 g/kg EtOH. A two-way mixed ANOVA found significant main effects of conditioning day [*F*(1.981, 97.08)=30.13, *p*<0.0001], and phenotype [*F*(2, 49)=93.26, *p*<0.0001], as well as a significant interaction between these two factors [*F*(4. 98)=70.87, *p*<0.0001] (***Figure 2F***). Post-hoc analyses found that both CTA-resistant and -sensitive rats consumed significantly less saccharin compared to saline-injected rats on conditioning days 2 [resistant: *q*(25.21)=12.71, *p*<0.0001; sensitive: *q*(19.6)=15.12, *p*<0.0001] and 3 [resistant: *q*(26.02)=15.75, *p*<0.0001; sensitive: *q*(34.97)=41.62, *p*<0.0001]. In addition, CTA-sensitive rats injected with 2.0 g/kg EtOH consumed significantly less saccharin than rats in the resistant phenotype on the second [*q*(22.19)=4.392, *p*=0.014] and third [*q*(15.71)=15.03, *p*<0.0001] conditioning days. Taken together, these data reveal two distinct phenotypes in male Long-Evans rats, with CTA-sensitive rats exhibiting significantly greater magnitude ethanol-induced CTA relative to CTA-resistant rats at each dose tested.

The same analysis in females produced results comparable to our observations in males. Specifically, in the 1.5 g/kg EtOH group, there was a significant main effect of phenotype [*F*(2, 47)=18.81, *p*<0.0001] and a conditioning day x phenotype interaction [*F*(4, 94)=19.30, *p*<0.0001], but no main effect of conditioning day (***Figure 3C***). Post-hoc analyses found significantly lower saccharin intake in both CTA-resistant [*q*(17.81)=5.154, *p*=0.005] and -sensitive phenotypes [*q*(16.02)=5.655, *p*=0.003] relative to saline on conditioning day 2. CTA-resistant females also consumed significantly less saccharin compared to saline-injected females on the third conditioning day [*q*(28.74)=5.562, *p*=0.001], as did CTA-sensitive females [*q*(30.47)=13.75, *p*<0.0001]. However, unlike in males injected with the same dose of ethanol, the difference in saccharin intake between CTA-sensitive and -resistant females did not emerge until the third conditioning day [*q*(22.97)=7.968, *p*<0.0001]. Comparisons between phenotypes in females injected with 2.0 g/kg EtOH uncovered significant main effects of conditioning day [(*F*(1.963, 94.20)=14.50, *p*<0.0001] and phenotype [*F*(2, 48)=49.24, p<0.0001] as well as a significant conditioning day x phenotype interaction [*F*(4, 96)=22.63, *p*<0.0001]. Post-hoc analyses found significantly lower saccharin intake in both phenotypes compared to saline controls on the second [resistant: *q*(17.79)=5.815, *p*=0.0018; sensitive: *q*(19.16)=10.49, *p*<0.0001] and third [resistant: *q*(33.84)=8.033, *p*<0.0001; sensitive: *q*(31.47)=19.51, *p*<0.0001] conditioning days. In contrast to females injected with 1.5 g/kg EtOH, saccharin intake was significantly lower in CTA-sensitive females injected with 2.0 g/kg EtOH compared to CTA-resistant females on both conditioning day 2 [*q*(22.98) = 3.82, *p* = 0.033] and 3 [*q*(15.31)=11.04, *p*<0.0001]. Taken together, these data indicate that, similar to males, female CTA-sensitive rats exhibit stronger ethanol-induced CTA than CTA-resistant rats. However, in contrast to males, the difference between CTA-sensitive and -resistant phenotypes in females injected with 1.5 g/kg EtOH was not evident until the third conditioning day.

### Dose and sex differences in the emergence of ethanol-induced CTA across phenotypes

To more explicitly examine whether the rate at which CTA sensitivity emerged differed across phenotypes and ethanol dose, we assessed within-group differences in the percent change in saccharin intake from baseline on conditioning days 2 and 3. We hypothesized that ethanol-induced CTA would emerge after fewer pairings in the 2.0 g/kg EtOH dose compared to the 1.5 g/kg EtOH dose in both male and female rats. In rats injected with 1.5 g/kg EtOH, one-sample t tests on conditioning day 2 revealed that saccharin intake was unchanged in CTA-resistant male and female rats (p>0.05). In contrast, CTA-sensitive rats exhibited a significant decrease in saccharin intake from baseline ([males: *t*(12)=4.888, *p*=0.0004; females *t*(12)=2.736, *p*=0.018] (**Figure 4A**). As expected, CTA-sensitive male and female rats continued to exhibit a reduce saccharin intake from baseline on the third conditioning day [males: *t*(12)=10.55, *p*<0.0001; females: *t*(12)=10.49, *p*<0.0001]. Interestingly, by the third conditioning day, CTA-resistant males exhibited a significant reduction in intake [*t*(12)=3.676, *p*=0.003], however, intake in CTA-resistant females still did not differ significantly from baseline levels [*p*>0.05] (***Figure 4B***). CTA-sensitive rats of both sexes injected with 2.0 g/kg EtOH exhibited a significant reduction in saccharin intake from baseline on conditioning day 2 [males: *t*(12)=7.2, *p*<0.0001; females: *t*(12)=3.274, *p*=0.007]. One-sample t tests also showed that CTA-resistant males had significantly reduced saccharin intake on conditioning day 2 after 2.0 g/kg EtOH [*t*(12)=3.671, *p*=0.003]. In contrast, CTA-resistant females injected with 2.0 g/kg EtOH did not reduce their saccharin intake on the second conditioning day [*p*>0.05] (***Figure 4C***). By the third conditioning day, CTA-resistant and -sensitive rats of both sexes that had saccharin paired with 2.0 g/kg EtOH exhibited a significant reduction in saccharin intake relative to baseline [CTA-resistant males: *t*(12)=2.839, *p*=0.01; CTA-resistant females: *t*(11)=3.166, *p=*0.009; CTA-sensitive males: *t*(12)=47.06, *p*<0.0001; CTA-sensitive females: [*t*(12)=25.98, *p*<0.0001] (***Figure 4D***). Altogether, these data uncover significant sex differences within the CTA-resistant phenotype with CTA-resistant females exhibiting greater resistance to ethanol-induced changes in saccharin intake than CTA-resistant males. This difference is eliminated after multiple pairings with high dose ethanol (2.0 g/kg) indicative of a dose-dependent increase in ethanol-induced CTA independent of phenotype.

**Figure 4.**
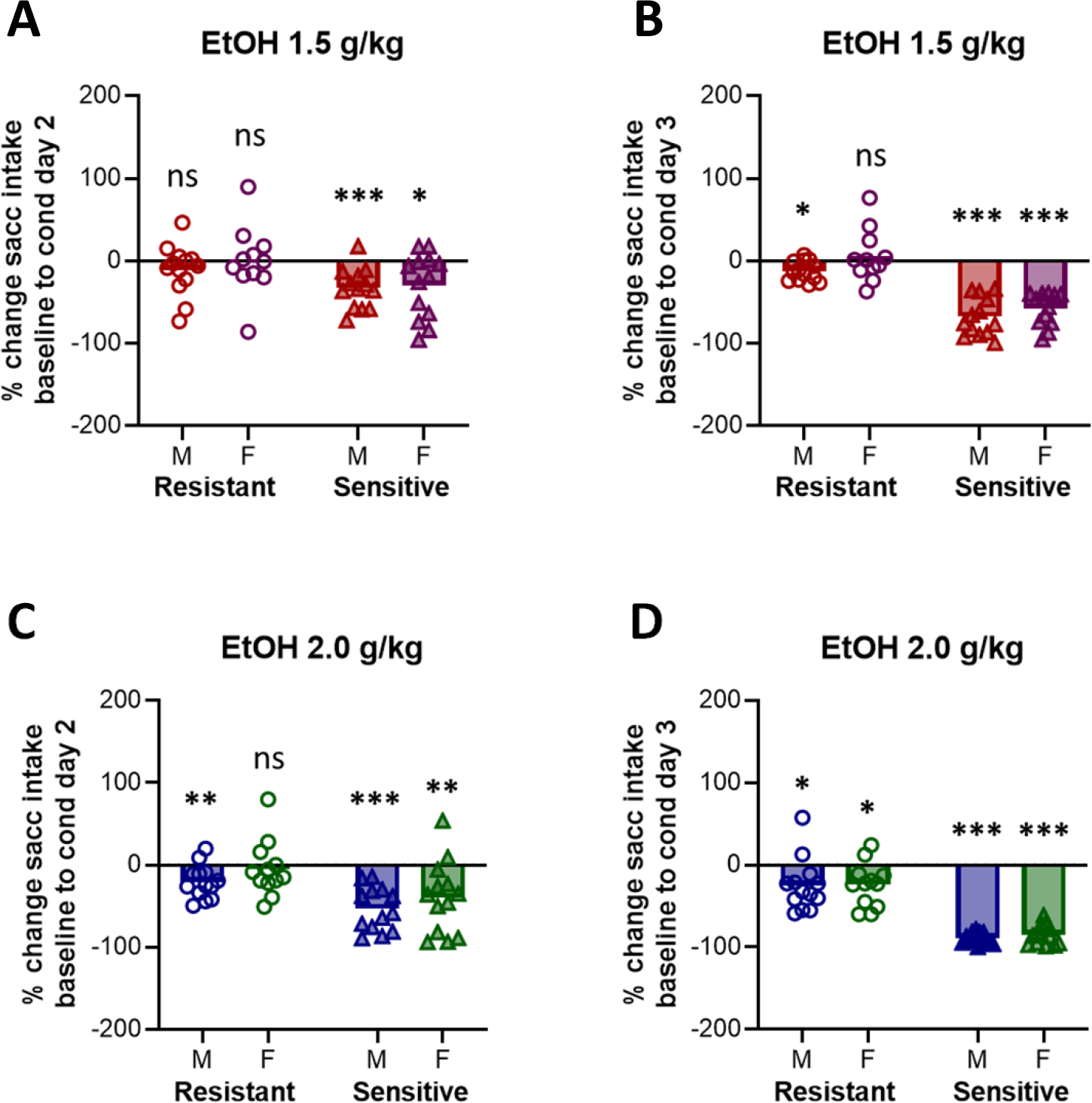
Phenotype-specific sex differences in CTA magnitude. **(A)** Comparisons of the percent change in saccharin intake from baseline across sexes within each phenotype revealed a significant decrease in intake on conditioning day 2 in CTA-sensitive male (M) and female (F) rats injected with 1.5 g/kg EtOH. In contrast, no significant change from baseline was observed in CTA-resistant rats of either sex. **(B)** By the third conditioning day, male CTA-resistant rats injected with 1.5 g/kg EtOH exhibited a significant reduction in saccharin intake from baseline levels, whereas saccharin intake in female CTA-resistant rats remained unchanged from baseline. **(C)** In the 2.0 g/kg EtOH group, CTA-resistant males exhibited a significant reduction in saccharin intake on conditioning day 2, whereas CTA-resistant females did not. **(D)** By the third conditioning day, both male and female CTA-resistant rats had significantly decreased their saccharin consumption relative to baseline levels, albeit to a lesser degree than CTA-sensitive rats. In contrast, CTA-sensitive rats of both sexes exhibited a significant decrease in saccharin intake on both conditioning days. Statistical comparisons to baseline: *p<0.05, **\*\***p<0.01, ***p<0.001, ns=not significant.

### Behavioral signs of intoxication across dose and sex

To determine whether behavioral signs of intoxication were distinct between CTA phenotypes, we first compared intoxication score between ethanol dose and phenotype averaged across all conditioning days. As expected, a two-way ANOVA revealed a significant main effect of ethanol dose [*F*(1, 48)=6.597, *p*=0.013] (***Figure 5A***), with males injected with 2.0 g/kg EtOH showing greater signs of intoxication compared to males injected with 1.5 g/kg EtOH. While CTA-sensitive rats appeared to have higher intoxication scores than CTA-resistant rats, this comparison did not reach statistical significance. To examine this more closely, we considered whether intoxication differed significantly between phenotypes during distinct phases of the CTA paradigm by comparing intoxication scores across conditioning days. A three-way ANOVA in males with conditioning day, dose, and phenotype as factors uncovered no significant main effects of any factor. There were also no significant interactions involving ethanol dose as a factor (*p*>0.05). Therefore, we performed a two-way ANOVA after collapsing data across doses. This analysis revealed a significant conditioning day x phenotype interaction [*F*(1, 38)=5.804, *p*=0.021] (***Figure 5B***). Post hoc analyses showed that CTA-sensitive males achieved greater intoxication scores after acute ethanol exposure during the first conditioning day compared to their CTA-resistant counterparts (*p*=0.006), although this difference did not persist on the second conditioning day (*p*>0.05). Given this difference between phenotypes on conditioning day 1, we next evaluated the relationship between level of intoxication on conditioning day 1 and CTA magnitude using a simple linear regression between intoxication score and percent change in saccharin intake. This analysis revealed a negative relationship between measures at both doses with greater intoxication scores associated with smaller percent change in saccharin intake. This relationship reached significance in the 2.0 g/kg group [*r^2^*=0.32, *p*=0.005] (***Figure 5D***), but not the 1.5 g/kg group [*r^2^*=0.041, *p*>0.05] (***Figure 5C***). These data suggest that the degree of ethanol-induced motor impairment experienced during first exposure to ethanol is predictive of CTA sensitivity – at least at higher doses of ethanol.

**Figure 5.**
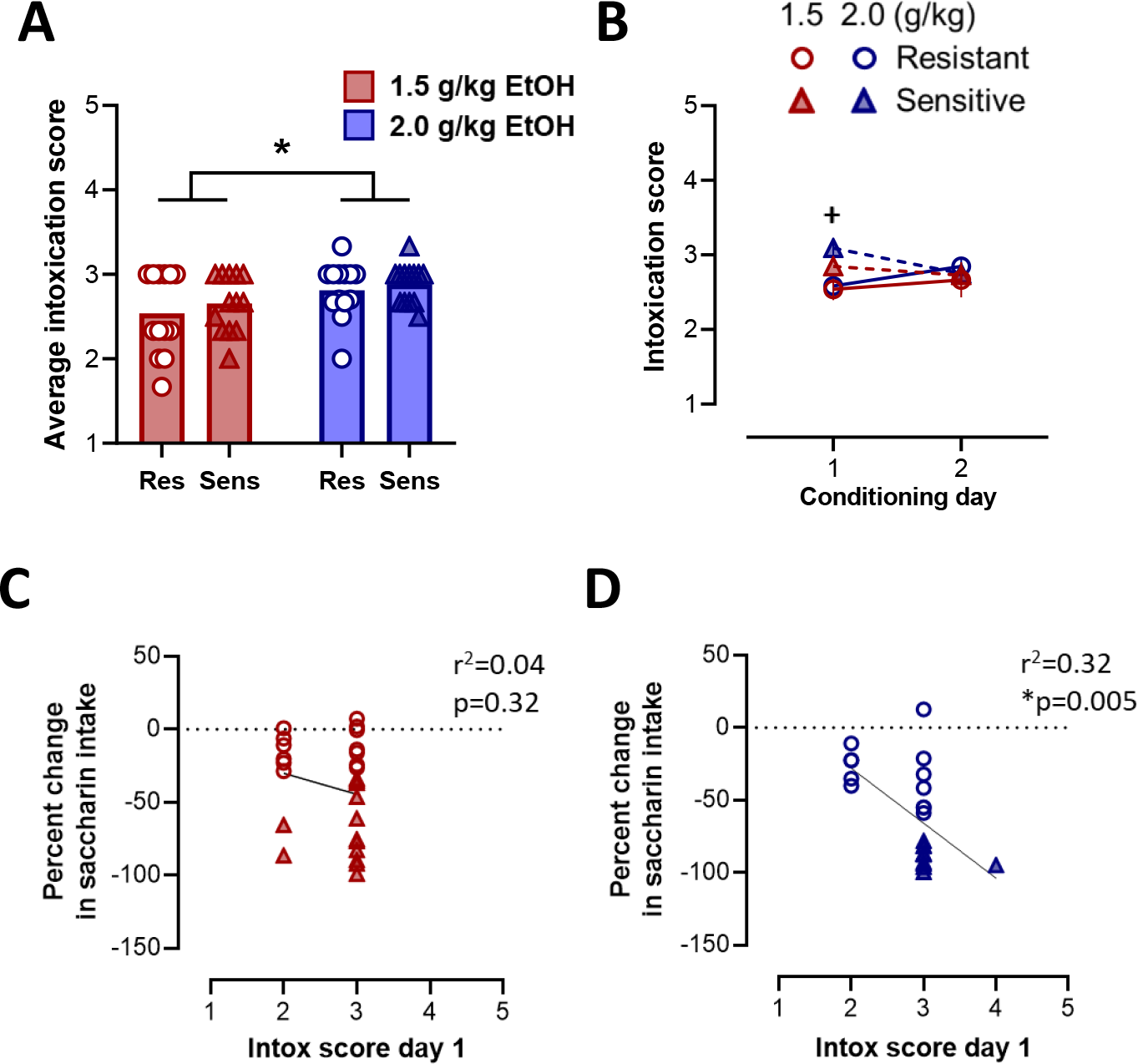
Relationship between behavioral signs of intoxication and CTA phenotype in males. **(A)** Rats injected with 2.0 g/kg EtOH achieved significantly higher intoxication scores across the entire CTA procedure than rats injected with 1.5 g/kg EtOH regardless of phenotype **(B)** When examined by conditioning day, CTA-sensitive rats achieved significantly higher intoxication scores on the first conditioning day compared to CTA-resistant rats at both doses of EtOH tested. **(C)** Intoxication score on the first conditioning day was not significantly correlated with CTA magnitude in rats injected with 1.5 g/kg EtOH. **(D)** However, a significant negative relationship between intoxication score on conditioning day 1 and percent change from baseline in saccharin intake on conditioning day 3 was observed in rats injected with 2.0 g/kg EtOH. *p<0.05 between EtOH doses; ^+^p<0.05 between phenotypes.

The same analysis of average intoxication score across conditioning days in females also revealed a main effect of dose [*F*(1, 45)=13.34, *p*=0.0007] (***Figure 6A***), with rats in the 2.0 g/kg EtOH group showing greater levels of intoxication than the 1.5 g/kg group. A three-way ANOVA assessing intoxication score across conditioning day, dose, and phenotype showed no main effects of conditioning day or phenotype and no interactions, but did reveal a main effect of ethanol dose [*F*(1, 86)=15.38, *p*=0.0002] (***Figure 6B***). Female rats injected with 2.0 g/kg EtOH showed greater signs of intoxication compared to rats injected with 1.5 g/kg EtOH, regardless of phenotype. Unlike our observations in males, there was no significant relationship between intoxication score on conditioning day 1 and percent change in baseline saccharin intake in females at either the 1.5 g/kg (*r^2^*=0.12, *p*>0.05) (***Figure 6C***) or 2.0 g/kg EtOH (*r^2^*=0.02, *p*>0.05) doses. These data indicate that, unlike in males, there is minimal relationship between CTA sensitivity and behavioral signs of intoxication after acute ethanol exposure in females.

**Figure 6.**
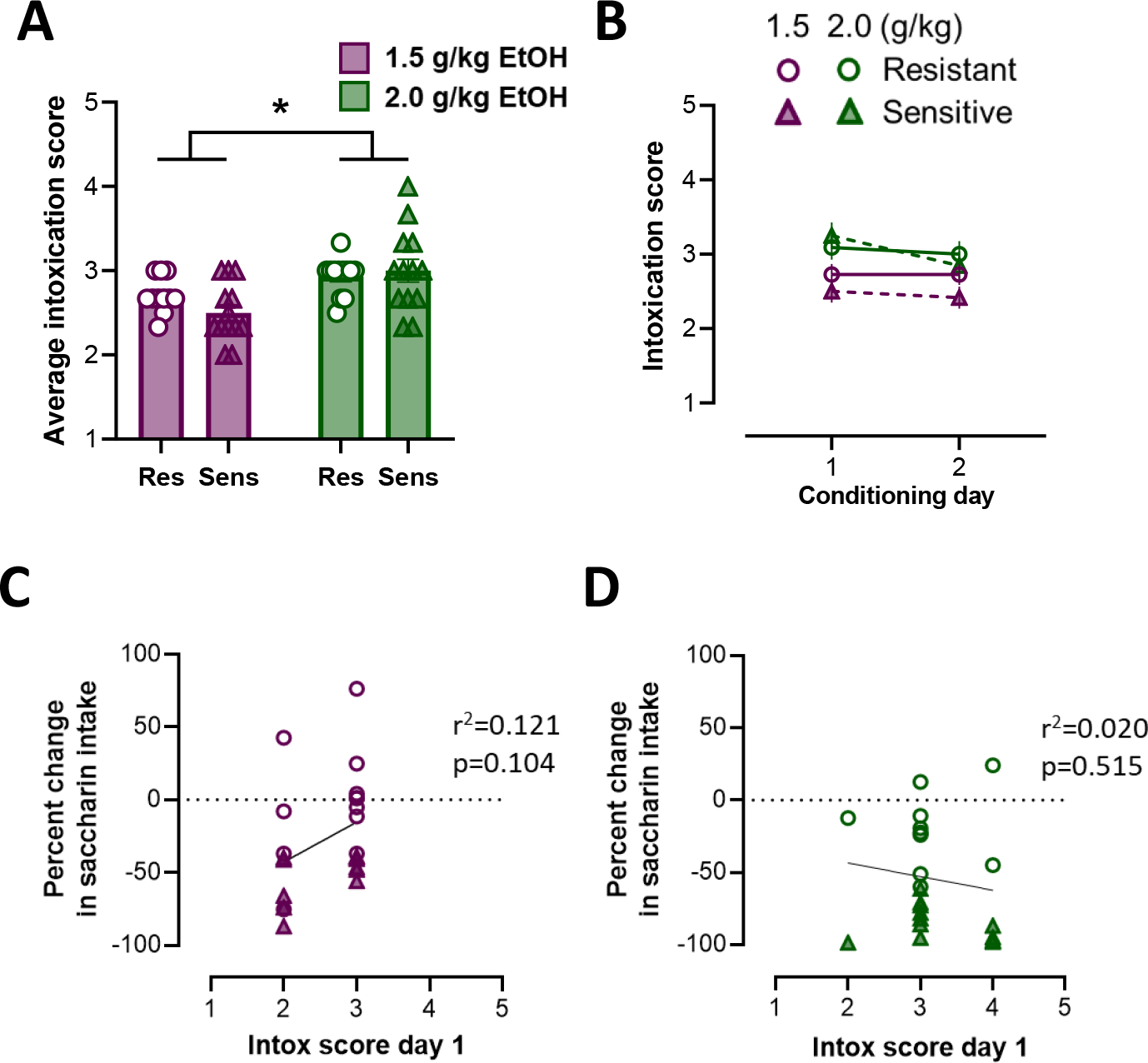
Relationship between behavioral signs of intoxication and CTA phenotype in females. **(A)** Rats injected with 2.0 g/kg EtOH achieved significantly higher intoxication scores compared to rats injected with 1.5 g/kg EtOH regardless of phenotype. **(B)** Unlike males, intoxication scores between female CTA-sensitive and -resistant rats were not different on either conditioning day. Intoxication score on conditioning day 1 was not significantly correlated with CTA induced by either 1.5 g/kg EtOH **(C)** or 2.0 g/kg EtOH **(D)**. *p<0.05 between EtOH doses.

## Discussion

Clinical studies have consistently found that individuals who are more responsive to the rewarding properties of ethanol and less responsive to its aversive properties are at higher risk for future binge drinking and AUD diagnosis (A. King et al., 2021; A. C. King et al., 2002, 2011, 2014, 2019b). While this relationship has been recapitulated in rodent models, preclinical findings have been limited to genetically and/or demographically distinct subpopulations (e.g., alcohol-preferring vs non-preferring; adult vs adolescent), unlike studies performed in clinical populations. Using CTA, results from the present study uncovered significant individual differences in response to the aversive properties of ethanol in adult male and female outbred Long-Evans rats. As expected, when examining group averages, the magnitude of ethanol-induced CTA was dose-dependent. However, closer inspection of individual responses revealed a bimodal distribution in CTA expression in both sexes at both doses. Thus, some individuals displayed a CTA-sensitive phenotype and others appeared relatively resistant to the development of ethanol-induced CTA. While the magnitude of ethanol-induced CTA was similar between male and female rats classified as CTA-sensitive, we observed significant sex differences in CTA-resistant rats, with females exhibiting greater resistance to the development of CTA than males. Altogether, these data uncover important individual differences in ethanol-induced CTA in a rodent model that more closely reproduces the variability observed in clinical samples than has been reported in previous preclinical studies.

We observed significant dose-dependent ethanol-induced CTA in adult male and female Long Evans rats, with 2.0 g/kg EtOH producing greater CTA magnitude than 1.5 g/kg EtOH in both sexes. These data are in agreement with prior research showing that ethanol becomes more aversive as dose increases (Cunningham, 2019; Moore et al., 2013; Morales et al., 2014; Phillips et al., 2005; Saalfield & Spear, 2016; Schramm-Sapyta et al., 2014; Vetter-O’Hagen et al., 2009b). Analysis of the distribution of CTA magnitude across individuals uncovered two distinct phenotypes present in both male and female rats: a CTA-sensitive group that exhibited the expected reduction in saccharin intake as the number of saccharin-ethanol pairings increased, and a CTA-resistant group that exhibited little-to-no reduction in saccharin intake across conditioning days. While both CTA-sensitive and -resistant phenotypes drank significantly less saccharin than saline controls over the course of conditioning, CTA-sensitive rats exhibited a rapid decrease in saccharin intake, reducing their intake by almost half after just one saccharin-ethanol pairing. This was followed by an even greater reduction in intake after the second saccharin-ethanol pairing. In contrast, CTA-resistant rats exhibited minimal reduction in saccharin intake after one saccharin-ethanol pairing. The majority of CTA-resistant rats then maintained this level of intake despite an additional conditioning day.

Notably, unlike males, CTA-resistant females that had saccharin paired with 1.5 g/kg ethanol exhibited a slight increase in saccharin intake over conditioning days, suggesting a near complete lack of sensitivity to the aversive properties of ethanol. This observation led us to more closely examine the rate at which rats within each phenotype developed CTA by assessing the percent change in saccharin intake from baseline on each conditioning day. These comparisons revealed that both male and female CTA-resistant rats maintained baseline levels of saccharin intake after a single pairing with 1.5 g/kg ethanol. While males reduced their intake significantly from baseline, albeit only slightly, female rats continued to maintain baseline levels of intake at this dose after an additional saccharin-ethanol pairing. Similar findings were observed in females after a single pairing with 2.0 g/kg ethanol, although this dose was sufficient to produce a slight, but significant, reduction in intake after two saccharin-ethanol pairings. In contrast, all CTA-sensitive rats exhibited a significant decrease in saccharin intake after a single saccharin-ethanol pairing regardless of sex or ethanol dose. Thus, our data indicate that sex differences in ethanol-induced CTA are driven primarily by females within the CTA-resistant phenotype. This distinction is particularly relevant given our inability to detect sex differences when data was averaged across the entire group (i.e., collapsed across phenotypes). Of note, conflicting results have been reported in prior studies with respect to the presence or absence of sex differences in ethanol-induced CTA. For example, some studies report similar ethanol- or lithium chloride-induced CTA between adult male and female rodents (Glover et al., 2016; Vetter-O’Hagen et al., 2009b). On the other hand, several studies have observed greater ethanol-induced CTA in males than females, although these differences depended on a number of variables including age, strain, and housing conditions (Morales et al., 2014; Schramm-Sapyta et al., 2014; Sherrill et al., 2011; Roma et al., 2006; Roma et al., 2007). Data from the present study suggest that differences in the prevalence of CTA-resistant and -sensitive phenotypes within a given sample may contribute to these contradictory results and that sex differences in CTA magnitude may be difficult to detect when examining population means. Instead, these differences may be better appreciated by comparing males and females within distinct phenotypes.

One possible explanation for the differences we observed in sensitivity to ethanol-induced CTA is that saccharin possesses greater hedonic value in CTA-resistant than -sensitive rats. Thus, it is plausible that in CTA-resistant rats the rewarding properties associated with saccharin consumption outweighed the aversive properties associated with acute ethanol exposure. This is unlikely, however, given that rats within a given sex exhibited similar levels of saccharin intake at baseline irrespective of phenotype or ethanol dose. It is also possible that the difference in CTA between resistant and sensitive phenotypes is not due to a difference in sensitivity to the aversive properties of ethanol, but rather reflects differential rates of learning between phenotypes. As such, CTA-resistant rats may require additional saccharin-ethanol pairings to more robustly associate saccharin consumption with the subjective effects of acute ethanol exposure. However, the slight, but significant, reduction in saccharin intake observed in most CTA-resistant rats on conditioning day 2 suggests that CTA-resistant rats were successful in making the association between saccharin consumption and acute ethanol exposure after a single pairing but simply did not find it sufficiently aversive to avoid saccharin to the same degree as CTA-sensitive rats. Moreover, data from a separate study in our lab found that the difference in saccharin consumption between phenotypes was maintained even with additional saccharin-ethanol pairings (Ramirez et al., 2023).

While differences in the level of intoxication reached could contribute to differences in CTA between phenotypes injected with the same dose of ethanol, previous work suggests that blood ethanol concentration (BEC) is not a reliable predictor of response to ethanol’s aversive properties (Roma et al., 2006, 2007; Sherrill et al., 2011). Although BECs were not measured in the current study, behavioral signs of intoxication are significantly positively correlated with BEC (Glover et al., 2019; Glover et al., 2021) and were assessed immediately after ethanol administration. As expected, we observed a dose-dependent increase in behavioral signs of intoxication in male and female rats of both phenotypes. Given that the measures of intoxication we obtained are largely reflective of signs of motor impairment and sedation, which are among the properties of ethanol considered aversive (Holdstock et al., 2000; A. C. King et al., 2002, 2011; Schuckit, 1994), we considered the possibility that rats with higher intoxication scores may also exhibit greater sensitivity to ethanol-induced CTA. In support of this, we observed a significant negative relationship between intoxication score and change in saccharin intake in males injected with 2.0 g/kg EtOH. This finding suggests that rats exhibiting the greatest degree of motor impairment after acute ethanol find ethanol to be more aversive when tested using CTA. However, this finding was only observed at one ethanol dose and only in males, limiting its generalizability. Of note, previous work has shown that female rats require higher BECs to exhibit the same behavioral signs of intoxication as male rats (Glover et al., 2021) suggesting that a higher dose of ethanol may be required to reveal the same relationship between motor impairment and ethanol-induced CTA in females. Thus, it is possible that measures of motor impairment and sedation can predict CTA magnitude at higher doses of ethanol, but additional work using more precise measures (e.g., rotarod) are needed to thoroughly investigate this possibility. Doing so in the context of the behavioral phenotypes uncovered in the current study may allow for the identification of additional indicators of sensitivity or resistance to the aversive properties of ethanol.

While survey-based studies can capture measures of rewarding and aversive effects of acute ethanol exposure in tandem in humans, separate assays are required to assess the full range of ethanol’s properties in rodents. Importantly, clinical work has shown that individuals who exhibit a low response to ethanol’s aversive properties while also exhibiting a high response to its rewarding properties are at greatest risk for heavy drinking and AUD diagnosis (A. King et al., 2021; A. C. King et al., 2011, 2014, 2019b). While the present study only measured the response to ethanol’s aversive properties, our data make clear the need to perform similar assessment of phenotypic differences in the response to ethanol’s rewarding properties. Related, a large body of work has characterized phenotypic differences in incentive salience toward cues predictive of rewards including ethanol. This work has shown that a subset of rats, referred to as sign-trackers, attribute incentive salience to reward-predictive cues, as evidenced by more time spent interacting with the cue. This is in contrast to goal-trackers, which respond more to the location of reinforcer delivery (Angelyn et al., 2021; Flagel et al., 2007; Flagel & Robinson, 2017; Robinson & Flagel, 2009; Valyear et al., 2017; Villaruel & Chaudhri, 2016). Importantly, the sign-tracking phenotype is associated with measures indicative of greater risk for relapse to ethanol and other drug seeking in both humans and rodent models (Cofresí et al., 2022; Colaizzi et al., 2020; Flagel et al., 2010; Robinson et al., 2014; Saunders et al., 2014). Interestingly, sign trackers are significantly more resistant to lithium chloride-induced reinforcer devaluation (i.e., CTA) than goal-trackers (Colaizzi et al., 2020; Kuhn et al., 2022). Although speculative, these data suggest the possibility that a subset of rodents exists that exhibit both a disproportionately high response to cues predictive of reward while also exhibiting resistance to devaluation with an aversive stimulus. Additional studies performing systematic and comprehensive phenotypic assessment of individual differences across multiple paradigms that assess both rewarding and aversive properties of ethanol will be crucial in determining the neurobiological mechanisms underlying such phenotypic differences.

In summary, the present study identifies innate differences in sensitivity to ethanol-induced CTA in outbred adult male and female Long-Evans rats. In doing so, our findings provide a much-needed animal model for studying the neural mechanisms underlying variability in subjective response to the aversive properties of ethanol. Using this model, future studies have the potential to provide crucial insight into the neurobiological factors that confer risk for heavy alcohol drinking and AUD.

## Acknowledgements

We would like to thank current and past members of the Glover Lab for their technical assistance including Christen Amegashie, Alex Brown, Nidhi Chetan, Kacey Clayton-Stiglbauer, Karl Bosque-Cordero, Andres Gascon, Shikun Hou, Nikki Kinarasri, Autumn Ollice, Arleen Perez Ayala, Jacqueline Sanchez, Shree Srinivasan, and Shannon Wheeler. This work was supported by NIH grants R01 AA029130 (EJG), P50 AA022538 (EJG), R00 AA024208 (EJG), T32 AA026577 (KRP).

